# Geometry of neural dynamics along the cortical attractor landscape reflects changes in attention

**DOI:** 10.1101/2025.08.08.669432

**Authors:** Hayoung Song, Ruiqi Chen, Thomas L. Botch, Todd S. Braver, Monica D. Rosenberg, Jeffrey M. Zacks, ShiNung Ching

## Abstract

The brain is a complex dynamical system whose activity reflects changes in internal states, such as attention. While prior work has shown that large-scale brain activity reflects attention, the mechanism governing this association in a time-varying and task-dependent manner remains unknown. Here, we tested a hypothesis that the geometry of neural dynamics on the attractor landscape, or the movement along the “hills and valleys”, reflects changes in attentional states over time and variations across controlled and naturalistic contexts. We fit a parametric dynamical systems model to fMRI data collected during rest, task performance, and naturalistic movie-watching. The model decomposes neural dynamics into components that are intrinsic versus extrinsically driven by stimuli. Model parameters were biologically meaningful, reflecting both cognitive states and individual differences. Model simulations revealed a set of attractors that mirrored functional brain networks, spanning the canonical gradient from sensorimotor to default mode network regions. The speed and direction of neural trajectories toward these attractors systematically varied across attentional states in a context-dependent manner. When participants were paying attention to effortful tasks, neural dynamics converged directly toward a task-relevant attractor, suggesting that it occupied a steeper region of the attractor landscape. In contrast, when participants were engaged in sitcom episodes, neural dynamics were in a flattened region of the landscape, directed away from the attractors. These findings demonstrate that while the positions of the attractors are largely determined by the cortical organization, the geometry of neural dynamics on the attractor landscape changes systematically across attentional states and situational contexts.

## Introduction

The brain is a multiscale system that dynamically evolves over time to produce behavior. Dynamical systems modeling, which formalizes how neural activity changes over time using differential equations, has played a central role in neuroscience (Breakspear, 2017; Vyas et al., 2020). Such models have uncovered numerous biophysical mechanisms of the brain, from the early work of Hodgkin and Huxley (1952) who described how ion currents generate action potentials, to Wilson and Cowan (1972) who described how excitatory-inhibitory neuronal interactions shape population-level dynamics.

More recent work has extended this approach beyond single cells and neuronal populations to characterize whole-brain dynamics (Honey et al., 2007; Deco and Jirsa, 2012; Demirtaş et al., 2019; Wang et al., 2019; Singh et al., 2020, 2025). From a dynamical systems perspective, large-scale brain activity unfolds over time as a trajectory within a high-dimensional state space, where each dimension typically represents the activity of a brain region. This state space is shaped by an attractor landscape, or hills and valleys that guide the trajectory of neural activity (**Figure 1**). Research has shown that trajectories visit a small number of recurring brain states, each defined by a unique pattern of regional activity and interactions (Baker et al., 2014; Chen et al., 2016; Vidaurre et al., 2017; Liu et al., 2018; Yousefi and Keilholz, 2021). These brain states function as attractors, or deep valleys in the landscape, where neural activity is most likely to converge (Kelso, 2012; Cocchi et al., 2017; Roberts et al., 2019; Chen et al., 2025). In turn, neural dynamics are characterized by transitions between or perturbations around these attractors.

**Figure 1.**
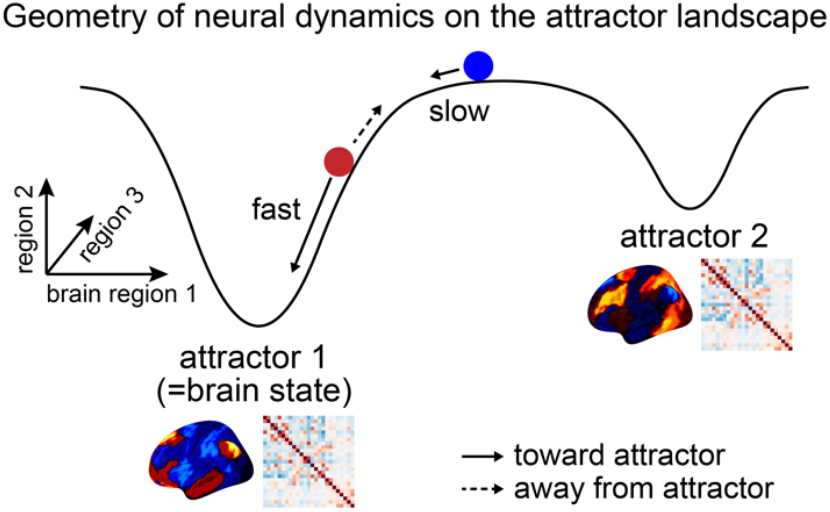
Schematics of the geometry of neural dynamics on the attractor landscape. A state space is defined where each dimension represents the activity of a brain region spanning the cortex. The hills and valleys represent the attractor landscape with valleys indicating the attractors. Each attractor corresponds to a recurring brain state that is identified from large-scale patterns of regional activity and interaction. The circles represent the neural activity at a specific moment. The trajectory of neural activity (indicated with black arrow) is largely determined by the landscape but can also be affected by external perturbations. For example, the red circle is more likely to fall toward the attractor based on the intrinsic landscape but may move away from the attractor when perturbed by external forces, such as stimuli, task demands, or behaviors. The speed and direction of the movement on this landscape defines the geometry of neural dynamics. Example brain state figures are adapted from Song et al. (2023).

While attractor landscapes govern neural dynamics, it is important to understand how they relate to our internal states, including physiological (e.g., hunger, stress, arousal) and cognitive states (e.g., attention, emotion, motivation) (Greene et al., 2023). Prior studies have provided initial evidence of this connection, showing that certain brain states are more or less likely to occur when a person is focused versus unfocused (Yamashita et al., 2021), performing better or worse on a task (Taghia et al., 2018; Cornblath et al., 2020), or engaged versus disengaged in a movie (Song et al., 2023). However, the computational mechanism of this connection remains unknown (John et al., 2022). We hypothesized that the geometry of neural dynamics on the attractor landscape—i.e., the speed and direction in which brain activity moves toward or away from the attractors—changes over time in relation to internal state fluctuations (**Figure 1**). Supporting this idea, Munn et al. (2021) showed evidence of flattened attractor landscape during phasic bursts in noradrenergic locus coeruleus activity (related to arousal; Sara, 2009) and steepened attractor landscape during phasic bursts in cholinergic basal forebrain activity (related to vigilance or attentional focus; Hasselmo and Sarter, 2011). This suggests a possibility that one’s internal state at a given moment may correspond to whether the brain occupies a flatter or steeper region of the attractor landscape, which in turn shapes the trajectory of neural dynamics.

Here, we propose that the geometry of neural dynamics on the attractor landscape characterizes moment-to-moment and context-to-context variations in internal states. In this study, we specifically test this in relation to measures of sustained attention. Dynamical systems models were fit to whole-brain fMRI data collected during rest, tasks, and movie-watching. Our model separates neural activity into two components: one that is intrinsic^1^, driven internally by regional activity and interactions, and the other extrinsic, driven externally from the stimulus. By simulating neural trajectories from the model, we identify attractors toward which the neural activity converges. At each moment, we estimate the speed and direction of these simulated trajectories, using them to infer the steepness of the local attractor landscape. Importantly, we relate the change in geometries to behavioral measures of attention which were collected with the fMRI data. These measures include participants’ continuous ratings of engagement while watching comedy sitcoms and button response times during controlled sustained attention tasks.

Prior work has typically fit a set of static model parameters from neural activity dynamics, meaning variations across time have often been reduced to a single model that is agnostic to temporal change. To the best of our knowledge, this is the first study to reproduce the evolving neural geometry from a model of fixed parameters and relate them to fluctuations in internal states. Using this novel approach, we test a hypothesis that the geometry of large-scale cortical dynamics along the attractor landscape systematically reflects changes in attentional states during task and movie-watching contexts.

### A dynamical systems model of large-scale cortical activity

We applied a large-scale parametric dynamical systems model, developed and validated by Singh et al. (2020) and Chen et al. (2025), to fit the time series of BOLD activity measured in human cortex with fMRI. The model defines the rate of change in neural activity 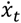, as a function of the neural activity *x*_*t*_ (*x*_*t*_ ∈ ℝ^*n*^, *n* = number of neural units) and time-aligned experimental variables *u*_*t*_ (*u*_*t*_ ∈ ℝ^*p*^, *p* = number of experimental variables) such as task, stimulus, or behavior. In our model, *x* corresponds to the BOLD activity time series of 200 parcels covering the cortex (Schaefer et al., 2018). *u* corresponds to audiovisual and semantic features extracted from the stimuli that participants watched and heard inside the scanner, which were reduced to 100 principal component dimensions. Mathematically, the model can be described as 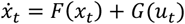, where *F* and *G* are deterministic functions that represent, respectively, the impact of intrinsic activity and input-driven perturbations on the evolution of neural activity.

The followings are the specifics of our model, where *F*(*x*_*t*_) = *Wψ*_*α*_(*x*_*t*_) – *D* ⊙ *x*_*t*_ and *G*(*u*_*t*_) = *βu*_*t*_ with *W, D, α*, and *β* being model parameters (**Figure 2**).

**Figure 2.**
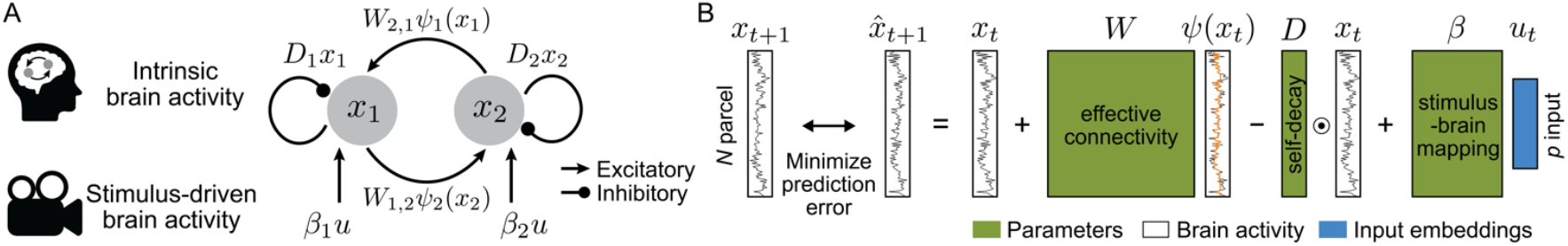
Model of large-scale cortical dynamics. **(A)** Model schematics. *x* represents the activity time series of a cortical parcel and *u* represents input from the stimulus. Model parameters include directional interactions between neural units (*W*), self-decay that determines autocorrelation (*D*), and the stimulus-to-brain relationship (*β*). Although only two units are visualized for simplicity, the model was fit on the time series of 200 cortical parcels. **(B)** Model optimization. The model was trained to minimize the difference between the observed and predicted neural activity patterns of consecutive time steps. Green denotes parameters that are estimated during training, black lines in denote observed neural activity pattern at a time step (*x*_*t*_), with orange indicating sigmoidal bound from -1 to 1 given the nonlinear transfer function, and blue denotes stimulus embeddings at the corresponding time step (*u*_*t*_).

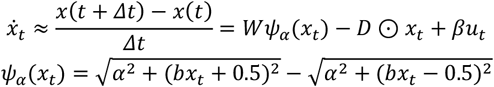

Given that *Δt* equals to 1 TR in our data, the equation can be simplified as follows.

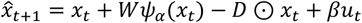

The goal of the model is to predict neural activity pattern of the consecutive time step 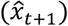, by minimizing the prediction error (i.e., difference between the predicted 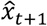 and observed *x*_*t*+1_) using algorithmic optimization (here, stochastic gradient descent). The prediction of the next time step is based on the neural activity *x*_*t*_ plus the signals received by connections from other parcels (*Wψ*_*α*_(*x*_*t*_)) minus the self-decay (*D* ⊙ *x*_*t*_) plus the neural activity driven by the external inputs (*βu*_*t*_). In turn, fitting this model corresponds to decomposing intrinsic and extrinsic (i.e., input-driven) neural dynamics.

The weight matrix (*W* ∈ ℝ^*n*×*n*^) represents directional interaction between neural units, namely the effective connectivity. Nonlinearity is introduced by a parametrized sigmoidal transfer function (*ψ*_*a*_), which maps neural activity to a bounded output at a range from -1 to 1. The slope of the transfer function differs for every parcel, parameterized by *α* (*α* ∈ ℝ^*n*^). *b* is fixed at 20/3 following prior studies (Singh et al., 2020; Chen et al., 2025). A self-decay (*D* ∈ ℝ^*n*^) captures a return to baseline in absence of external inputs or interactions, with ⊙ denoting element-wise product. Having high decay rate corresponds to having low temporal autocorrelation, indicating less persistence in neural activity. *β* represents linear transformation from the stimulus embedding space to neural activity pattern space. Note that this is an individualized model where parameters are estimated for each fMRI run of each participant.

We analyzed two openly available fMRI datasets, the SONG dataset (N=27; Song et al., 2023) and the HCP dataset (N=119; Barch et al., 2013; Van Essen et al., 2013; Finn and Bandettini, 2021). In both datasets, each participant underwent multiple sessions of rest, task, and movie-watching conditions. We extracted time series of stimulus features for the movie watching runs, including low-level visual and visuo-semantic features of the video frames, low-level audio of the sounds, and audio-semantic features of the speech and dialogues in the movies. These features were projected onto the 100 principal components that explained the largest variance across runs. Only low-level visual and visuo-semantic features were extracted from the SONG dataset task runs because the stimuli were purely visual. Given that no stimulus was provided for resting-state runs in either dataset, we fit a simplified model that only considers intrinsic neural dynamics but not extrinsic neural dynamics: 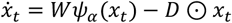 (removing the input term *βu*_*t*_ from the equation).

### Model parameters recapitulate functional brain connectivity and stimulus encoding

We first tested whether our model, optimized to predict neural activity of successive time steps, captured parameters sensitive to individual and cognitive state differences. These properties are critical, as their emergence indicates the model’s biological plausibility. We analyzed the movie-watching runs of the SONG and HCP datasets because they were fit on the full model that includes the input term.

Model performance was estimated based on how well the model predicted neural activity of the next time steps, specifically 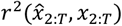. The total explained variance was, on average, *r*^2^ = 0.452 ± 0.043 across a total of 80 runs in the SONG dataset and *r*^2^ = 0.561 ± 0.053 across 476 runs in the HCP dataset (**Figure 2A**). This indicates that our model explained approximately half of the variance in neural activity. We then decomposed the explained variance into three components. The interareal interaction explained *r*^2^ = 0.041 ± 0.009 (SONG), 0.055 ± 0.015 (HCP), local recurrence explained *r*^2^ = 0.284 ± 0.051, 0.237 ± 0.060, and external input-driven activity explained *r*^2^ = 0.017 ± 0.003, 0.015 ± 0.004 of the variances.

We further validated the model by comparing model parameters, *W* and *β*, to their analogue descriptive statistics that are commonly used in the field. *W* was compared to an undirected functional connectivity (FC), estimated as the parcel-by-parcel Fisher’s transformed Pearson’s correlation coefficients. *β* was compared to the regression coefficients estimated from a linear encoding model, that predicts neural activity from stimulus time series: *x*_*t*_ = (encoding coefficient) × *u*_*t*_ + (residual) (encoding coefficient ∈ ℝ^*n*×*p*^, residual ∈ ℝ^*n*^). We found that the estimated *W* was highly comparable to FC for both the SONG (cosine similarity = 0.929 ± 0.009; *z* = 1178.36, *p* < .0001 compared to shuffled chance distribution) and HCP datasets (cosine similarity = 0.913 ± 0.014; *z* = 2830.05, *p* < .0001) (**Figure 3B, D**). Likewise, *β* was highly comparable to encoding coefficients for both the SONG (cosine similarity = 0.324 ± 0.100; *z* = 448.58, *p* < .0001) and HCP datasets (cosine similarity = 0.243 ± 0.036; *z* = 1022.18, *p* < .0001) (**Figure 3C**).

**Figure 3.**
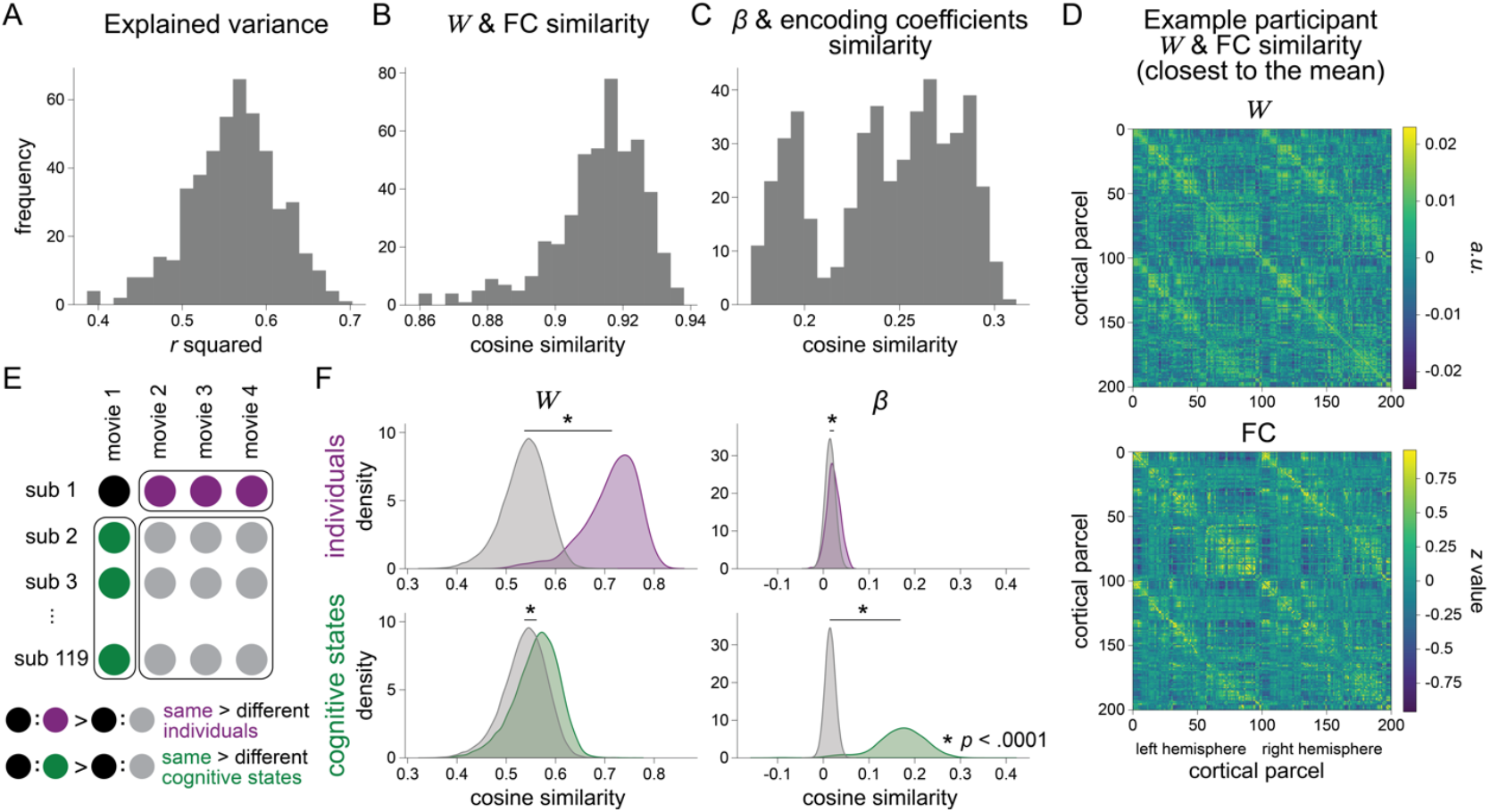
Model validation with the HCP dataset. **(A)** Model performance was primarily assessed based on the explained variance of how well the model predicted neural activity of the next time step given the current time step (the explicit training objective). The histogram includes estimates from all movie-watching runs of 119 participants included in the HCP dataset. **(B-C)** Model parameters were compared to descriptive statistics, specifically the similarities of *W* to functional connectivity (FC) estimates and *β* to coefficients estimated by the stimulus-to-brain encoding models. These aspects of the model were not explicitly optimized. **(D)** Example participant’s *W* estimate compared to their FC matrix. A representative participant’s data was selected for visualization (whose cosine similarity between *W* and FC was closest to the mean in **B**). **(E)** Individual differences were assessed by comparing parameter similarities between different movie watching runs of the same participant (black-purple) to those of different participants (black-grey). Cognitive state differences were assessed by comparing parameter similarities of different participants’ same movie-watching runs (black-green) to those of different movie-watching runs (black-grey). **(F)** Density functions comparing similarities in parameter estimates between the same vs. different individuals (*top*) and the same vs. different cognitive states elicited by the same vs. different movies (*bottom*). Lines on top of the density functions connect the means of the two distributions, with asterisks indicating statistical significance. **Supplementary Figure S1** shows the same model validation results, analyzed with the SONG dataset.

Do these parameters capture differences between individuals as well as differences in cognitive states? Models were considered sensitive to individual differences if the parameters estimated from runs of the same person were more similar compared to parameters of different individuals (**Figure 3E**). Both *W* and *β* were sensitive to individual differences, and *W* more strongly reflected individual differences than *β* (Wilcoxon rank-sum test between same vs. different individual pairs; *W*: *z* = 14.957, *p* = 1.4e-50 for SONG, *z* = 44.667, *p* = 0.0 for HCP; *β*: *z* = 2.714, *p* = 0.007 for SONG, *z* = 13.828, *p* = 1.7e-43 for HCP) (**Figure 3F**). This aligns with prior findings that functional connectivity is stable within an individual and distinctive across individuals, thus driven more by trait than state differences (Cole et al., 2014; Gratton et al., 2018).

Models were considered sensitive to cognitive state differences if parameters estimated from runs of the same movie stimulus were more similar compared to runs of different stimuli (**Figure 3E**). Both *W* and *β* were sensitive to cognitive state differences, and *β* more strongly reflected cognitive state differences than *W* (*W*: *z* = 21.414, *p* = 9.9e-102 for SONG, *z* = 75.509, *p* = 0.0 for HCP; *β*: *z* = 45.189, *p* = 0.0 for SONG, *z* = 242.537, *p* = 0.0 for HCP) (**Figure 3F**). The corresponding descriptive statistics—the FC and encoding coefficients—closely followed this trend. These results indicate that the parameters estimated from our dynamical systems model are comparable to validated descriptive statistics and hold biological plausibility.

### The model reveals stable attractors organized along the cortical hierarchy

In dynamical systems, the differential equation determines and predicts how the system’s trajectory would evolve over time from a given initial state, assuming that no internal or external perturbation exists beyond what is parameterized. Depending on the nature of the system, the trajectory may eventually converge to a set of stable fixed point attractors, settle onto stable periodic orbits known as limit cycles, bounce between saddle points when attractors are absent, or exhibit chaotic behavior (Chen et al., 2025).

We hypothesized that our data-driven models would reveal a finite set of attractors, or stable patterns of neural activity where neural trajectories are more likely to converge. To find attractors, we ran forward simulations of the model starting at various initial states in the observed fMRI time series (i.e., each initial state was an observed 200-parcel neural activity pattern at a chosen time step) (**Table 1**). We simulated

**Table 1.**
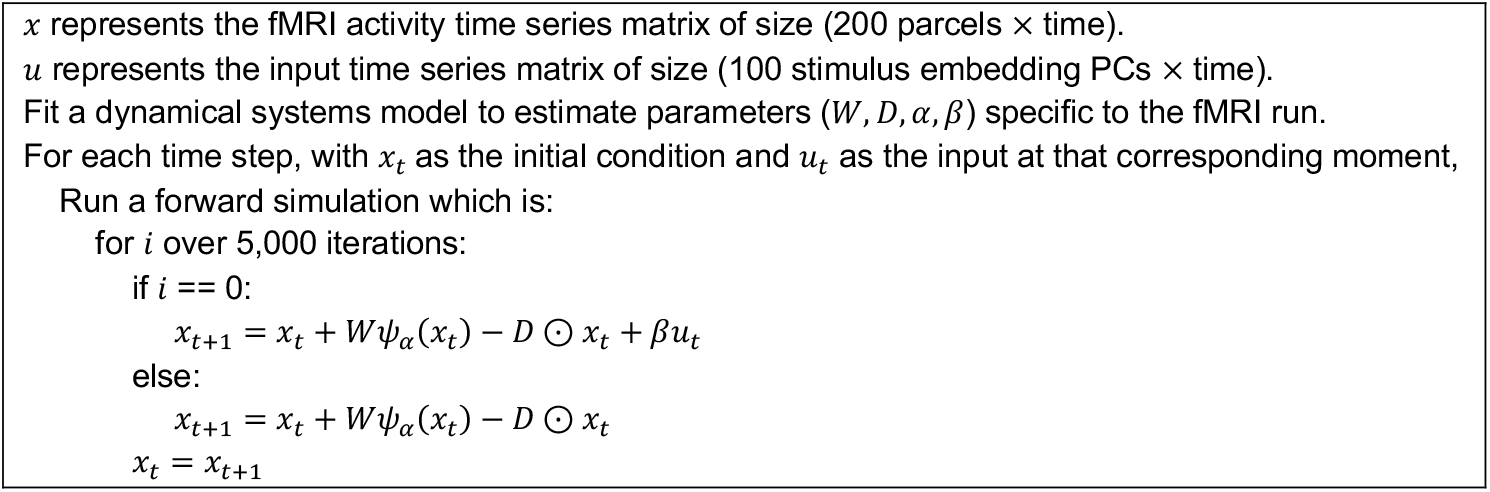
Forward simulation. For every time step of the neural data, we simulated the model continuously to predict the likely path of neural dynamics, assuming it strictly follows the equation of the model. For each time step, *βu*_*t*_ was simulated only one time step in the future because stimulus-driven activity for future simulations cannot be predicted. In contrast, the intrinsic drift could be predicted through simulations, by using the predicted *x*_*t*+1_ as the next *x*_*t*_ and iterating this 5,000 times. The attractors identified this way represent states that the neural activity tend toward, assuming no further perturbation beyond what’s driven by the input *u*_*t*_.

5,000 timesteps forward in time, which we considered was sufficient for convergence. This method simulates the intrinsic drift, or the likely path that neural trajectories will take, assuming no perturbation to the drift other than what is parameterized. That is, it allowed us to infer the attractor landscape of large-scale cortical dynamics.

We focused this analysis on the HCP dataset, given that it contained a larger number of total runs (1,666 runs) compared to the SONG dataset (188 runs). We included rest, task, and movie-watching runs in the analysis. When conducting forward simulations on all runs, a majority of runs converged onto a set of point attractors: many converged onto 2 (41.06%) or 4 attractors (51.08%), and some converged onto 6 (7.02%) or 8 attractors (0.66%) (**Figure 4A**). Only 3 out of 1,666 runs (0.18%) did not converge to fixed point attractors but exhibited oscillatory limit cycles. This indicates that when large-scale cortical activity evolves according to the dynamics specified by the model equations, it is most likely to fall into a set of attractors. Note that the model algorithm is designed to identify an even number of attractors, each representing opposing patterns of neural activity.

**Figure 4.**
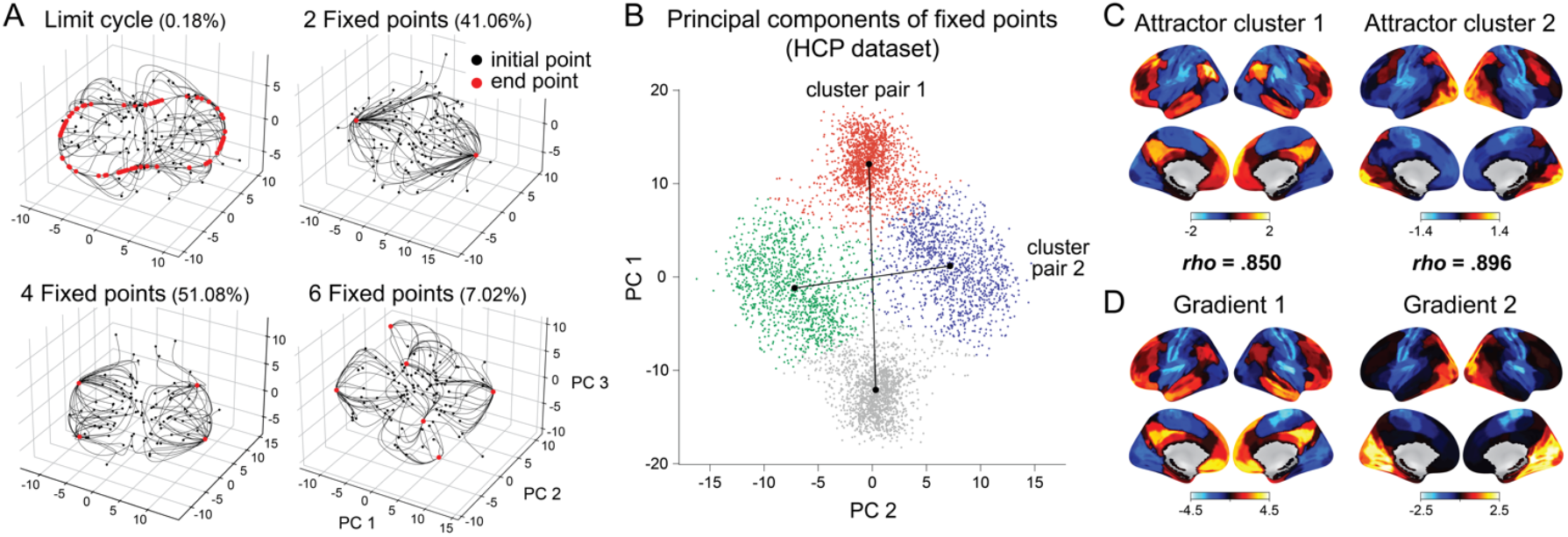
Attractor landscapes of large-scale cortical activity. **(A)** Four types of forward simulation results visualized from example fMRI runs. Black dots indicate 100 randomly sampled initial states (i.e., 100 time steps sampled from the observed neural activity). Each line indicates a trajectory taken over the course of a forward simulation (5,000 time steps). Red dots indicate the end states of the simulations, corresponding to the attractors. Principal component analysis was conducted per run to visualize the results in a 3D principal component (PC) space. **(B)** Attractors identified from all runs of the HCP dataset, projected onto the shared PC space and color-coded based on the outcome of k-means clustering into 4 clusters. Dots correspond to attractors estimated from the 1,666 HCP runs. Lines connect the pairs of cluster centroids that have anticorrelated patterns. **(C)** Neural patterns of the two identified attractor clusters. **(D)** Neural patterns of the top two gradients identified from Margulies et al. (2016). *ρ* values in between **C** and **D** indicate rank correlation values of the top and bottom neural patterns.

We predicted that these attractors would tile the core gradients of cortical organization that spans between the transmodal default mode network areas and the unimodal sensory and motor areas (Mesulam, 1998; Margulies et al., 2016). Furthermore, we expected that these attractors would correspond to canonical brain states—distinct and recurring patterns of neural activity—that have been replicated in multiple studies (Bolt et al., 2022; Song et al., 2023). To test these hypotheses, we combined the activity patterns of all attractors estimated in every run. We applied a k-means clustering to find 4 attractor clusters, because a majority of runs exhibited either four or less attractors (**Figure 4B**).

The mean activity patterns of these attractor clusters (**Figure 4C**) were compared to the known cortical gradients estimated by Margulies et al. (2016)^2^ (**Figure 4D**). Our two attractor clusters were highly comparable to the top cortical gradients (*ρ* values = .850 and .896, *p* values < .0001). Specifically, the attractor clusters served as axes that separated i) the default mode network (DMN) from the unimodal (UNI) sensory and motor areas and ii) the visual network (VIS) from the sensorimotor network (SM). These results provide a dynamical systems explanation for the recurrence of a small number of brain states over time: brain states correspond to attractors, the positions of these attractors are constrained by the brain’s functional network organization, and neural dynamics unfold along this attractor landscape.

### Geometry of neural dynamics differs across fluctuating attentional states during task and movie-watching

We characterized the attractor landscapes of large-scale neural dynamics, or the likely path that neural activity will take when following the model’s equations (**Figure 4A**). Would the geometry of these paths vary depending on a person’s attentional state at a given moment? In other words, would one’s attentional state—from a state of being focused on a task versus zoning out to being engaged in a movie versus being bored—relate to where the brain state is positioned within the hills and valleys?

Revisiting the forward simulation in **Table 1**, we identified two vectors at each time step based on the observed neural activity *x*_*t*_: one defining the intrinsic drift, or the flow governed by regional connections and self-decays (called the “intrinsic vector” defined by *Wψ*_*α*_(*x*_*t*_) ™ *D* ⊙ *x*_*t*_, shortened as 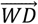), and the other defining the flow nudged by the external inputs at the corresponding moment (called the “extrinsic vector” defined by *βu*_*t*_, shortened as 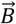). To quantify this, for every time step, we extracted the intrinsic and extrinsic vectors and calculated their angles and magnitudes (**Figure 5A**). The angle indicates the direction of the vector with respect to the position of the attractor it eventually converged, with high angle indicating the vector directing away from the attractor. The magnitude represents the degree of change from the initial state, with high magnitude indicating a fast-moving vector. Because these measures were estimated at each time step, we were able to extract their time series over the course of the fMRI run. In essence, they provide a geometric characterization of neural dynamics on the attractor landscape.

**Figure 5.**
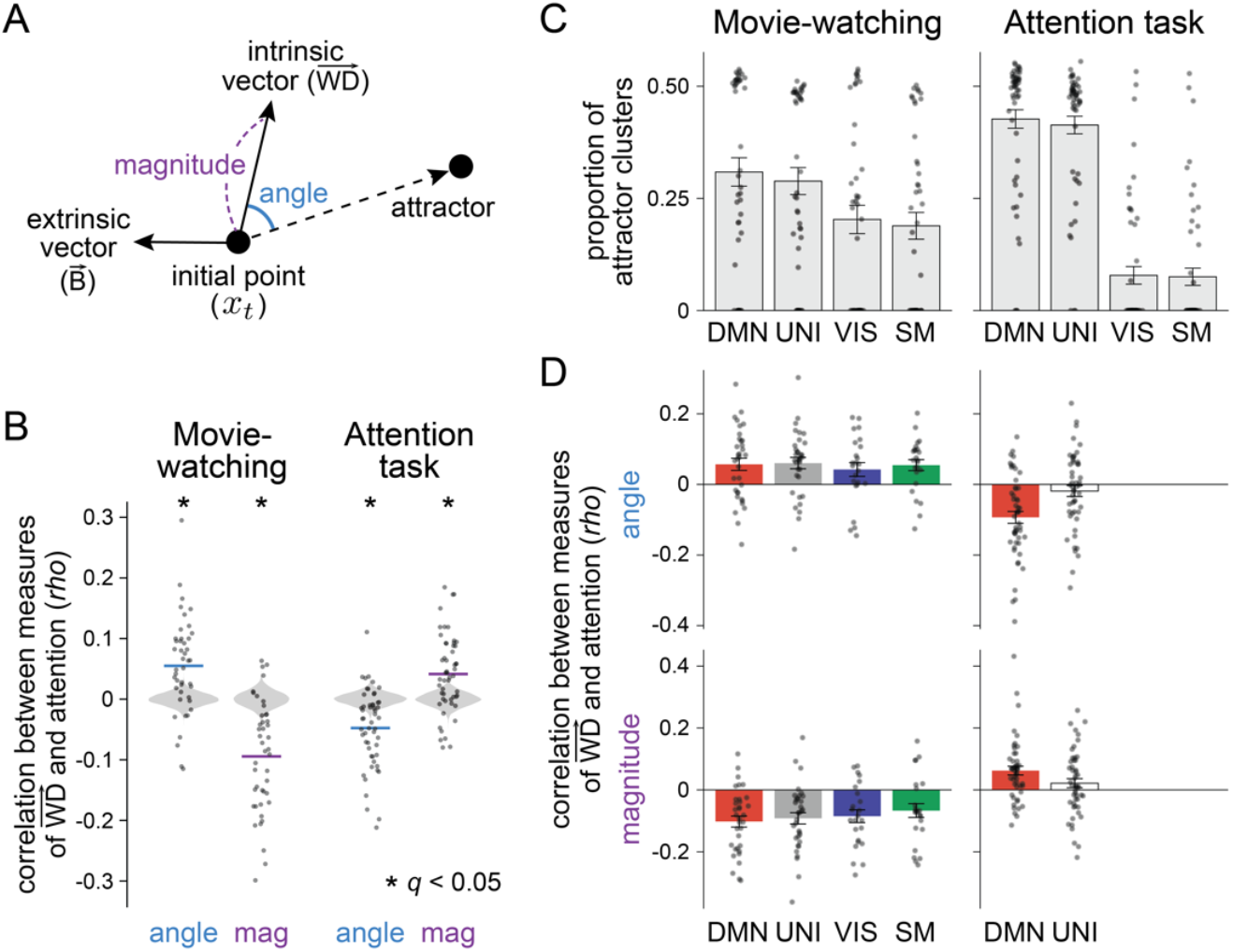
Neural dynamics toward attractors and their relationship with attention. **(A)** Geometric measures were estimated from neural activity at every time step at the first instance of forward simulation. In *x*_*t*+1_ = *x*_*t*_ + *Wψ*_*α*_(*x*_*t*_) ™ *D* ⊙ *x*_*t*_ + *βu*_*t*_, we decomposed a vector representing the intrinsic drift (*Wψ*_*α*_(*x*_*t*_) ™ *D* ⊙ *x*_*t*_) and a vector driven by external inputs (*βu*_*t*_). The magnitude of the vector was calculated using the Euclidean norm. The direction of the vector was calculated using the cosine angle with respect to the position of the attractor. Because these four measures were calculated at every time step, we were able to extract their respective time series for each fMRI run. **(B)** Correlations between attention measures and the angle (blue) and magnitude (purple) of the intrinsic vectors across movie-watching and attention task runs, compared to the respective chance distribution. Light grey areas indicate permuted chance distributions, black dots indicate correlation values between attention measures and the estimated angle or magnitude, and colored lines indicate the mean of these correlation values. Asterisks denote statistical significance shown in **Table 2.** Mag: magnitude. **(C)** Proportions of identified attractor clusters. The identified attractor at each time step was categorized to either one of the two ends of the primary gradient (default mode network [DMN] or unimodal network [UNI]) or the two ends of the secondary gradient (visual network [VIS] or somatosensory-motor network [SM]). The proportion of the attractor cluster was calculated at each run (black dot), which was averaged across runs to be summarized as a bar graph. Error bar indicates the standard error of the mean. **(D)** Correlations between individuals’ attention measures and the angle and magnitude of the intrinsic vector, categorized based on the positions of the attractors. Black dots indicate estimates from every run of every participant. Colored bars indicate the mean of correlation values that are significantly different from the permuted chance distribution (corrected for false discovery rate, *q* < .05), whereas empty bars indicate non-significance. Because VIS and SM attractors were less likely to occur during tasks, they were excluded from the analyses. **Supplementary Figure S3** shows separate results for the two runs in each context.

We asked whether neural trajectories toward attractors—characterized by the angle and magnitude of the intrinsic and extrinsic activity—systematically varied based on the attentional state of a participant. We focused our analysis on the SONG dataset, which contained both naturalistic movie-watching (i.e., two runs of comedy sitcom episode watching) and controlled attention tasks (i.e., two runs of gradual-onset continuous performance task; gradCPT) (Song et al., 2023). Importantly, these runs included time-varying behavioral measures of attention. During sitcom episodes, participants continuously rated how engaging they found the episode by adjusting the scale bar (Song et al., 2021).^3^ This measure was collected after the fMRI scan, as participants re-watched the same episode. During gradCPT, participants pressed a button at every second whenever target images appeared inside the scanner. The inverse of response time variability served as a proxy of sustained attention, with moments of stable response times indicating high attention and variable response times indicating low attention (Rosenberg et al., 2013). We correlated the behavioral time series of each participant with the four geometric measures’ time series, which was compared to a respective chance distribution where each geometric measure was correlated with circular-shifted behavioral time courses (**Table 2**; **Figure 5B**).

**Table 2.** Correlations between attention measures and the angle (representing direction) and magnitude (representing speed) of the neural trajectory across different conditions, compared to the respective chance distribution. False discovery rate (FDR) correction was applied to control for multiple comparisons across 16 significance tests. Asterisks indicate FDR-corrected *p* < .05.

|  | Movie |  | Task |  |
| --- | --- | --- | --- | --- |
|  | sitcom ep 1 | sitcom ep2 | gradCPT face | gradCPT scene |
| angle ($\overrightarrow{WD}$, attractor) | $z=3.570, q=0.002^*$ | $z=4.047, q<.0001^*$ | $z=-3.451, q=0.002^*$ | $z=-3.277, q=0.002^*$ |
| magnitude ($\overrightarrow{WD}$) | $z=-4.779, q<.0001^*$ | $z=-5.776, q<.0001^*$ | $z=2.147, q=0.048^*$ | $z=3.096, q=0.005^*$ |
| angle ($\vec{B}$, attractor) | $z=-2.275, q=0.040^*$ | $z=0.773, q=0.440$ | $z=1.817, q=0.093$ | $z=1.815, q=0.093$ |
| magnitude ($\vec{B}$) | $z=-4.778, q<.0001^*$ | $z=-1.377, q=0.207$ | $z=1.137, q=0.293$ | $z=0.825, q=0.432$ |

**Table 3.**
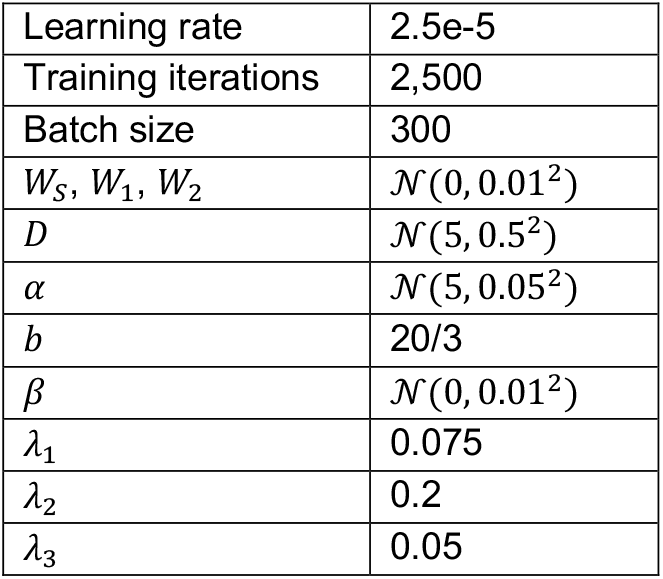
List of hyperparameters and parameter initializations.

|  |  |
| --- | --- |
| Learning rate | 2.5e-5 |
| Training iterations | 2,500 |
| Batch size | 300 |
| $W_S, W_1, W_2$ | $\mathcal{N}(0, 0.01^2)$ |
| $D$ | $\mathcal{N}(5, 0.5^2)$ |
| $\alpha$ | $\mathcal{N}(5, 0.05^2)$ |
| $b$ | 20/3 |
| $\beta$ | $\mathcal{N}(0, 0.01^2)$ |
| $\lambda_1$ | 0.075 |
| $\lambda_2$ | 0.2 |
| $\lambda_3$ | 0.05 |

Both the angle and magnitude of intrinsic activity 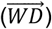 were significantly correlated with attention dynamics (**Table 2**; **Figure 5B**). The relationships were comparable between repeated runs of the movie-watching and attention tasks, highlighting the reliability of our results. Interestingly, an opposite relationship to attention dynamics was found between the two contexts. The angle of the intrinsic vector was large when participants reported high engagement toward episodes, whereas the angle was small when participants performed stably in gradCPT. The magnitude of the intrinsic vector was small when participants reported high engagement toward episodes, whereas the magnitude was large when participants performed stably in gradCPT. This means that neural dynamics toward attractors—largely determined by the landscape—not only varied across attentional state dynamics but in a manner different across contexts.

On the other hand, the angle and magnitude of the extrinsic activity 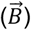 did not meaningfully relate to attention (**Table 2**). There were weak trends of correlations, which were seemingly derived from emergent correlations amongst the four geometric measures (**Supplementary Figure S2**). This highlights that attention relates to changes in intrinsic neural dynamics, but not stimulus-driven neural dynamics.

In a study by Song et al. (2023) in which this original data was collected, different brain states were associated with high attention in movie-watching and attention task contexts. Participants reported high engagement to sitcom episodes during the “base” state, a state where no functional networks exhibited dominant activity and was positioned at the center of the latent manifold. On the contrary, participants exhibited stable task performance to gradCPT during the DMN state, a state defined by high activity in the DMN (consistent with findings by Esterman et al., 2014; Fortenbaugh et al., 2018; Kucyi et al., 2020).

Motivated by these results, we categorized attractors into either of the four ends of gradients 1 (DMN and UNI) and 2 (VIS and SM), based on the neural pattern similarity between attractors and the gradients. We assessed the relationships between the geometric measures and attention dynamics, separately for these four attractors. The goal was to see if the relationship, shown in **Figure 5B** and **Table 2**, differs depending on where the neural trajectories converge.

We primarily found that the likelihood of attractors differed across the two contexts (**Figure 5C**). The neural activity was nearly equally likely to fall into one of the four attractors during movie-watching, whereas the ends of gradient 1, the DMN and UNI, were much more likely to serve as attractors during sustained attention tasks. This finding showing different likelihood of attractors across contexts indicates that the attractor landscapes differed across contexts.

During movie-watching, we found that the main relationship between the geometric measures and attention dynamics remained consistent, irrespective of which attractors the neural activity fell into. When participants reported high engagement to movies, the intrinsic drift directed away from the attractors with decreased magnitude (**Figure 5D**). This implies that the brain was in a flattened or shallow region of the landscape when people were attentive, such that the brain activity was more likely to lie at the center of the manifold and gravitated less toward the attractors (**Figure 6A**). On the contrary, during attention tasks, the main effect we found in **Table 2** was specific to when the neural activity converged onto the DMN attractor (**Figure 5C**). When attentive, neural activity approached the DMN attractor faster and more directly. This implies that the brain was on a steeper region of the attractor landscape near the DMN attractor when attentive (**Figure 6B**).

**Figure 6.**
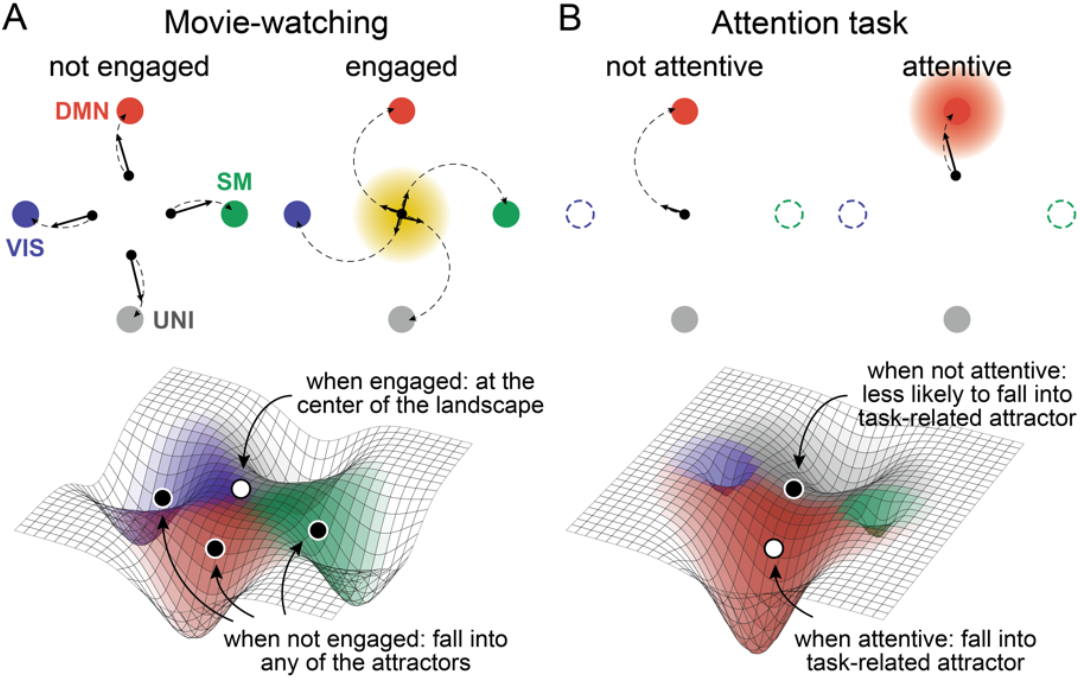
Schematic illustration of the results in **Figure 5D.** (*Top*) The angle and magnitude of the intrinsic vector during engaged vs. not engaged and attentive vs. not attentive states in movie-watching and attention task contexts. The black node illustrates the initial state and the four colored nodes in the surrounding illustrate attractors, with the vertical axis indicating gradient 1 and the horizontal axis indicating gradient 2 as in **Figure 4B**. The illustration is based on the angle and magnitude of the vectors from the initial state to the attractors, not the positions of the initial states with respect to the attractors nor the distance amongst attractors. Black dashed lines illustrate intrinsic drifts of neural activity toward attractors. Black solid lines illustrate a one-time-step vector of the drift. Shaded areas indicate positions of brain states that were reported to be associated with high attention in Song et al. (2023). The VIS and SM attractors are dashed in attention task context because they are less likely to occur in this context (**Figure 5C**). (*Bottom*) The 3D schematic illustrations of the attractor landscapes in the two contexts. White circles indicate brain states in attentive states and black circles indicate brain states in inattentive states.

Together, the results suggest that the neural dynamics toward attractors differed across high and low attentional states, in a context-dependent manner. This indicates that the attractor landscape of large-scale cortical dynamics systematically varies depending on attentional states and task demands.

## Discussion

In this study, we fit a dynamical systems model to large-scale fMRI data to investigate how the geometry of neural dynamics differs across changes in attentional states in different contexts. By simulating brain dynamics over time, we identified stable attractors that aligned with cortical gradients separating transmodal from sensorimotor regions as well as sensory from motor regions. Neural trajectories toward these attractors systematically varied with moment-to-moment attentional state fluctuations, in a manner different across situational contexts. These results indicate that the geometry of neural dynamics along the attractor landscape reflects changing attentional states and different task demands across situations.

The dynamical systems model used in this study is a simplified neural mass model that is tailored to simulate large-scale regional activities and their interactions, rather than local neuronal activities within a region. We found that the model parameters effectively reproduced descriptive statistics such as functional connectivity and stimulus encoding and were sensitive to trait- and state-level differences. Beyond reproducing descriptive statistics, the model’s strength comes from decomposing components of neural activity that are driven by interactions between brain regions, autocorrelation within each region, and external inputs—which together explained nearly half of the variance in the observed neural activity. These parameter estimates became the basis for estimating attractors and trajectories along the landscape. Moreover, the model was not tailored specifically to fit fMRI data, meaning the model can be generalized to other data modalities. These together suggest that our model provides a mathematical description of neural dynamics that are generative, biologically plausible, and generalizable to other research domains.

Forward simulation of the model revealed a set of attractors to which neural activity was more likely to converge. These attractors recapitulated canonical brain states that were identified in previous studies, which tiled the known gradients of cortical hierarchy. Specifically, the attractors were marked by high activities in the default mode network, visual network, and somatosensory-motor network— an emergent property of the model rather than a feature imposed by the model design. This conceptually replicates many studies in human systems neuroscience that have revealed the existence of a low-dimensional manifold of macroscale neural activity (Margulies et al., 2016; Hong et al., 2020; Shafiei et al., 2020; Dong et al., 2021) that is conserved across evolution (Oligschläger et al., 2019; Xu et al., 2020), stable across contexts (Cross et al., 2021; Samara et al., 2023), and confined by structural architecture and genetic makeup of the brain (Burt et al., 2018; Paquola et al., 2019; Vázquez-Rodríguez et al., 2019; Pang et al., 2023). This indicates that the positions of the attractors are largely fixed, confined by the brain’s canonical functional architecture.

Although the positions of the attractors are largely determined, it does not mean that the attractor landscape is fixed. Rather, the attractor landscape as well as the traversal along its hills and valleys can be flexible, which motivated us to study neural dynamics along the attractor landscape in relation to attention dynamics. We found that not only did the attractor landscapes differ across situational contexts, but even within a context, regions occupied within the landscape varied depending on a person’s attentional state. When participants were engaged in sitcom episodes, the intrinsic drift directed away from the attractors with decreased magnitude. This was a depiction of neural activity being less prone to fall into attractors as it situated on a shallow landscape at moments of engagement. In contrast, when participants were paying attention to an effortful psychological task, the intrinsic drift directed specifically toward the DMN attractor with increased magnitude. This indicates that the neural activity more easily fell into the DMN attractor because it was in a steeper landscape when attentive. This is in line with studies that associated high activity in DMN with moments of optimal performance during this task (Esterman et al., 2014; Fortenbaugh et al., 2018; Kucyi et al., 2020). These results highlight the flexible geometry of neural dynamics on the large-scale attractor landscape—it systematically changes across attentional state fluctuations.

The findings that brain activity tended to lie on a shallow attractor landscape when engaged in sitcoms and a steep local attractor when attentive to tasks resemble results reported by Munn et al. (2021). This past work found evidence of flattened cortical landscape upon activation of the noradrenergic arousal system, which projects broadly across the cortex, and deepened cortical landscape upon activation of the cholinergic vigilance system, which projects to relatively local functional networks. This suggests a hypothesis that different neuromodulatory circuits may underlie internal states of being immersed in engaging narratives versus being attentive to effortful and controlled tasks. In line with previous findings that engagement correlates with perceived emotional arousal during narratives (Busselle and Bilandzic, 2009; Bilandzic et al., 2019; Song et al., 2021; Ke et al., 2025), our results hint that the attractor landscape of being engaged may resemble a state of heightened arousal, more so than heightened vigilance. Future work can address this hypothesis by administering pharmacological agents that modulate noradrenergic and cholinergic activity during similar task and movie-watching conditions.

In sum, the attractor landscape is flexible across situational contexts, and cortical dynamics along this landscape reflect changes in attentional states. By modeling neural dynamics, we offer a new framework that can explain how internal states arise from large-scale brain activity.

## Acknowledgment

We thank JeongJun Park with conceptualization and helpful discussions and comments on the manuscript. The research was supported by the McDonnell Center for Systems Neuroscience and the McDonnell Center for Cellular and Molecular Neurobiology at Washington University in St. Louis (HS).

## Data and code availability

Model and analysis codes and human fMRI and behavioral data are openly available in: https://github.com/hyssong/dynamicalsystems

## Methods

### Model description

The dynamical systems model used in this study is adopted from the neural mass model called the mesoscale individualized neurodynamic (MINDy) model (Singh et al., 2020; Chen et al., 2025). The model is designed to fit the neural activity time series—whichever units or scales the neural activities are sampled from—and the time-aligned experimental variables, such as task, stimulus, or behavior. For our use, the neural activity corresponded to the BOLD activity of 200 cortical parcels collected from human fMRI. The experimental variable corresponded to audiovisual and semantic feature embeddings of the movies that participants watched inside the scanner and visual feature embeddings of the images that were presented as task stimuli. However, the choice of neural units and experimental variables can vary depending on the study and research question. No tailoring specific to the fMRI data (e.g., deconvolution of the hemodynamic response function) was made.

To reiterate, the model is defined as the following equation, with the neural activity at time *t* represented as *x*_*t*_ (*x*_*t*_ ∈ ℝ^*n*^, *n* = 200 parcels) and stimuli at time *t* represented as *u*_*t*_ (*u*_*t*_ ∈ ℝ^*p*^, *p* = 100 PCs of the feature embeddings), with *Δt* set to 1 TR.

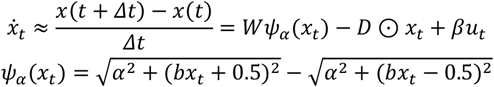

The weight matrix (*W* ∈ ℝ^*n*×*n*^) represents directional interaction between neural units, which conceptually corresponds to effective connectivity. *Wψ*_*α*_(*x*_*t*_) represents the weighted sum of the neural unit’s nonlinearly transformed activity at time *t* multiplied by its directed connections from every other neural unit. To constrain the estimation of *W*, we decomposed the parameter into *W* = *W*_*S*_ + *W*_*L*_ where *W*_*S*_ (*W*_*S*_ ∈ ℝ^*n*×*n*^) represents a sparse matrix after *L*_1_ regularization and 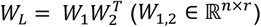 is given by low-rank approximation (*r* = *n*/3) with sparsity also given to *W*_1_ and *W*_2_ with *L*_1_ regularization.

Nonlinearity follows a parametrized sigmoidal transfer function (*ψ*_*a*_), which maps neural activity to a bounded output at a range from -1 to 1. The slope of the transfer function is determined by the estimated *α. b* is fixed at 20/3 in our model.

*β* ∈ ℝ^*n*×*p*^ represents a linear coupling between neural activity and incoming stimuli that are time-aligned with one another. The time was aligned by convolving the stimulus time series with the canonical hemodynamic response function. *β* conceptually corresponds to a linear mapping from the stimulus space to the neural space.

A decay term (*D* ∈ ℝ^*n*^) represents convergence to baseline activity at the absence of external inputs or interactions. *D* is an algorithmically important parameter, because *D* is initially set to a value significantly higher than 1 which flips the right-hand side of the equation toward a large negative factor of *x*_*t*_. This accentuates the difference in neural activity between units thus allowing effective estimation of the directed connectivity *W*. From our empirical tests, if *D* is set to a biologically plausible value of < 1, the estimated *W* does not recapitulate the descriptive functional connectivity measure.

### Model fitting

We fit the neural activity time series acquired at each run, in batches of size 300 TRs. Specifically, we predicted the neural activity of consecutive time steps 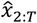 based on the observed neural activity *x*_1:*T*–1_ (where *T* = 300). Model parameters were optimized across 2,500 iterations using stochastic gradient descent, specifically the Nesterov-accelerated Adaptive Moment Estimation (NADAM) optimizer as chosen in previous studies (Singh et al., 2020; Chen et al., 2025). The loss ℒ was calculated as follows. *Λ* represents the regularization term which enforces sparsity in connections. Regularization terms were fixed to *λ*_1_ = 0.075, *λ*_2_ = 0.2, *λ*_3_ = 0.05.

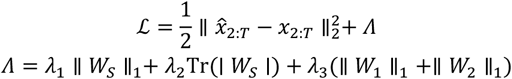

Because *D* was initialized at a value higher than 1, the prediction accuracy (i.e., rank correlation between the predicted and observed neural activity patterns) started at a value near -1 at the start of the training. The prediction accuracy increased gradually across iterations. In contrast, *W* quickly became comparable to a functional connectivity matrix in the initial phase of training, but across more iterations, it gradually approached a diagonally dominant matrix, which is conceptually similar to the first-order autoregressive model. To prioritize biologically meaningful parameter optimization rather than brain activity prediction, we stopped the training at 2,500 iterations (in batches of 300 consecutive TRs) at which point *W* was similar to functional connectivity but the model’s accuracy still remained negative. To boost prediction accuracy, we fit an ordinary least squares linear regression model to estimate *pW, pD*, and *pB*, which are scalar values (*pW, pD, pB* ∈ ℝ). Parameters *W, D*, and *β* were scaled by these value estimates respectively.

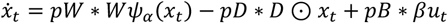

### Hyperparameters

Hyperparameter selection largely followed the original implementation of the MINDy model (Singh et al., 2020), but with some simplifications made.

### FMRI datasets

Two openly available fMRI datasets were analyzed: the SONG dataset (participants recruited in South Korea; N=27) and the HCP dataset (participants recruited in the USA; N=119), with preprocessing steps following Song et al. (2023). We applied a 200-parcel cortical atlas by Schaefer et al. (2018) where the BOLD activities of voxels corresponding to each parcel were averaged to represent the parcel activity (Singh et al., 2020; Chen et al., 2025). The SONG dataset includes two runs of resting-state, two runs of controlled sustained attention task called the gradual-onset continuous performance task (gradCPT), two runs of comedy sitcom-watching, and one run of educational documentary-watching (3T scans with TR = 1s). The HCP dataset includes four runs of resting-state in 3T (TR = 0.72 s) and four in 7T (TR = 1 s), two runs of working memory tasks in 3T (6 different types of cognitive task runs were excluded from analyses because the total TRs were less than 350 TRs), and four runs of movie-watching in 7T. These runs varied in total duration, ranging from 405 to 1486 TRs.

### Input features

Low-level visual features were characterized by hue, saturation, and pixel intensity, which were estimated per frame and averaged across frames within an event (rgb2hsv function in MATLAB). In the gradCPT run with face images, hue and saturation were excluded from the analyses because the images were presented in grayscale. Low-level audio features were represented with amplitude and pitch of left and right stereos, which were estimated per frame and averaged across frames within an event (audioread and pitch functions in MATLAB). Visuo-semantic features were represented by 512-dimensional embeddings of OpenAI’s pretrained Contrastive Language-Image Pre-training (CLIP) model (Radford et al., 2021) (huggingface; clip-vit-base-patch32). For audio-semantic features, we first applied OpenAI’s WhisperX model to the audio file to transcribe speech and dialogues, along with their time stamps (Bain et al., 2023). Transcripts were sampled at a TR resolution and were transformed using 512-dimensional embeddings of Google’s Universal Sentence Encoder (USE) (Cer et al., 2018) (tensorflow; https://tfhub.dev/google/universal-sentence-encoder-multilingual/3). A mean value of the respective dimension was assigned to moments when speech or dialogue did not exist.

Embedding time series were convolved with a canonical hemodynamic response function and normalized across time per dimension. Principal component analyses were conducted on the concatenated embedding time series of the 3 movie-watching runs in SONG and 4 runs in HCP respectively. We selected the top 100 principal component time series, which explained 74.97% and 77.69% of variance respectively. Principal component analysis was conducted on the gradCPT face run in isolation, given that the same images were presented in the same sequence for all participants (100.00% of explained variance for 100 principal components). For gradCPT scene runs, because presented images and sequences differed for all participants, participant-specific embeddings were concatenated for a principal component analysis (85.34% of explained variance). Resulting principal component time series were again normalized across time per dimension to be used as stimulus input *u* in the model.

### Validation of model parameters

Parameters were estimated through training the model on the data collected from an individual fMRI run (2,500 training iterations). After parameters were fixed, we predicted the next-time-step neural activity from the observed neural activity time series to calculate explained variance (*r*^2^). Note that the data used for training and testing were the same. When comparing parameter *W* with an undirected functional connectivity matrix (parcel-by-parcel Fisher’s transformed Pearson’s correlation coefficients), we took the average of the upper and lower triangles of *W* (region *i* → *j* and region *j* → *i* in a directed graph) and took the cosine similarity with the edge strengths of the functional connectivity matrix. Encoding coefficients were estimated using an ordinary least squares regression with a residual term. Cosine similarity between all values in *β* and values in encoding coefficients were computed. For both metrics, values in descriptive statistics were randomly shuffled 10,000 times which served as the respective chance distribution. *Z* statistics and two-tailed *p* values were calculated with respect to chance distributions. Cosine similarities were calculated between parameter estimates of run pairs. The cosine similarity values were grouped into whether they correspond to the same vs. different individuals or the same vs. different movie stimuli, which were compared using Wilcoxon rank sum tests.

### Attractor clusters and gradients

Attractors were estimated by forward simulating the observed neural activity pattern at each time step based on the estimated model parameters (**Table 1**). For each time step, we performed 5,000 simulations and recorded the resulting neural activity pattern, referred to as the “end state”. If the Euclidean distance between end states from different time steps was less than 0.1, those were grouped as the same attractor. Using this approach, we found that most fMRI runs converged to 2 or 4 attractors. Principal component analysis was applied to each individual fMRI run for visualization in a 3D space (**Figure 4A**).

We then aggregated attractors (each defined as a neural activity pattern across 200 parcels) across all fMRI runs and applied k-means clustering to identify 4 attractor clusters. The choice of k = 4 was motivated by the observation that most runs exhibited either 2 or 4 attractors, with higher numbers occurring less frequently. The mean activity pattern of each cluster represented the attractor cluster (**Figure 4B** and **4C**). Two pairs of clusters exhibited a pattern correlation of -1. Principal component analysis was also applied to the aggregated attractor patterns for visualization (**Figure 4B**).

Cortical voxel gradients estimated by Margulies et al. (2016) were obtained from NeuroVault (https://identifiers.org/neurovault.collection: 1598). Gradient values were averaged within each parcel to generate gradient estimates of the 200 parcels (**Figure 4D**). The neural activity patterns of the attractor clusters were then compared to the parcel-level gradient values using Spearman’s rank correlation.

### Relating speed and direction of neural dynamics toward attractors to measures of attention

With the neural activity pattern at each time step as an initial state, we considered one-step forward simulation of *x*_*t*+1_ = *x*_*t*_ + *Wψ*_*α*_(*x*_*t*_) – *D* ⊙ *x*_*t*_ + *βu*_*t*_ and separated vectors representing internally-driven change (*Wψ*_*α*_(*x*_*t*_) – *D* ⊙ *x*_*t*_, shortened to 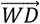) and externally-driven change (*βu*_*t*_, shortened to 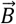). The angle between the respective vector and the eventual end state was calculated with the following equation,

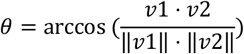

where *v*1 · *v*2 represents the dot product of the two vectors, and ‖*v*1‖ and ‖*v*2‖ are the Euclidean norms of the vectors. Before applying the inverse cosine, the cosine similarity was clipped to the range of [-1, 1]. The magnitude of the vector was calculated as the Euclidean norm. Because these measures were estimated at each time step, repeating this across all time steps within a run resulted their time series.

To probe attentional state changes during movie-watching, we asked participants to continuously rate their engagement, on a scale of 1 to 9, as they re-watched the sitcom episodes after the fMRI scan. Each participant’s engagement rating was normalized across time and convolved with a hemodynamic response function. To probe attentional state changes during gradCPT, we analyzed participants’ button response times. After linear interpolation of no button response trials and regressing out the linear trend, we calculated response time variability by taking the deviance from the mean response time at every TR. Because studies showed that moments of low response time variability correspond to high sustained attention or optimal task performance, the inverse response time variability time course was used as a proxy for attention dynamics (Rosenberg et al., 2013). Again, the inverse response time variability was normalized across time and convolved with a hemodynamic response function. See Song et al. (2023) for details of behavioral experiments and data analyses.

Angle and magnitude time series were correlated with attention measures sampled at a TR resolution using Spearman’s rank correlation, for each individual run. The mean of all participants’ Fisher’s *r*-to-*z* transformed *ρ* values were compared to permuted null distributions to calculate *z* statistics, where behavioral time series were circular-shifted across time with a random multiplication of either 1 or -1 (10,000 iterations). FDR correction was applied across 16 significance tests.

The relationship between geometric measures and attention was analyzed separately depending on the position of the attractor at each time step, or where the eventual end state lies at the final round of forward simulation. Because we found that attractor clusters lie at the ends of the known primary and secondary gradients (**Figure 4**), we categorized the attractor of each time step to one of the four ends of the two gradients. The attractor was labeled as either the DMN, UNI, VIS, or SM, based on the highest correlation coefficient. Correlation between the geometric measure and attention was calculated from a subset of time series corresponding to the respective attractor cluster category. In a majority of participants’ gradCPT runs, VIS and SM attractors did not appear. Because fewer than 10 fMRI participants’ gradCPT face or gradCPT scene runs included VIS and SM attractors, statistical analysis was not conducted for these cases. Significance was again tested by comparing the mean of Fisher’s *r*-to-*z* transformed *ρ* values to null distributions created from circular-shifted behavioral time series. Twelve comparisons were corrected for in **Figure 5D**, and 24 comparisons were corrected for in **Supplementary Figure S3B** using FDR correction.

## Supplementary Information

### Song *et al*. Geometry of neural dynamics along the cortical attractor landscape reflects changes in attention

**Supplementary Figure S1.**
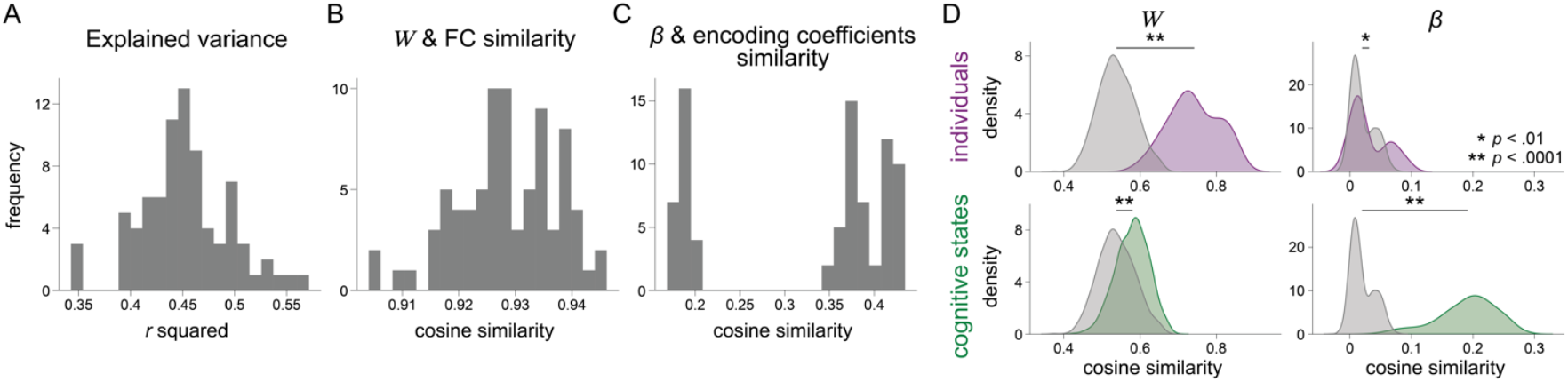
Model validation with the SONG dataset. The figure complements **Figure 3**.

**Supplementary Figure S2.**
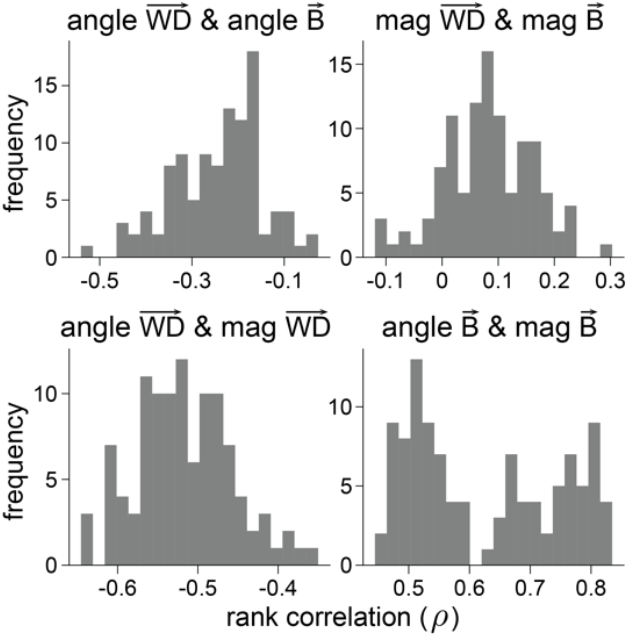
Correlations between geometric measures estimated from forward simulation. We asked whether the angle and magnitude of the intrinsic and extrinsic vectors were correlated with or independent from one another. Figures show that they are significantly correlated with one another. Time series of these measures were correlated in pairs in each run, and values across movie-watching and attention task runs in the SONG dataset comprised the histograms. At moments when the angle between 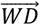 and the attractor was large, it was more likely that the angle between 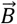 and the attractor was small. The angle and magnitude of 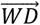 were negatively correlated, whereas the angle and magnitude of 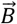 were positively correlated with one another. This dependence was not expected by the algorithm, meaning it was an emergent property upon fitting our model to the fMRI data. Mag: magnitude.

**Supplementary Figure S3.**
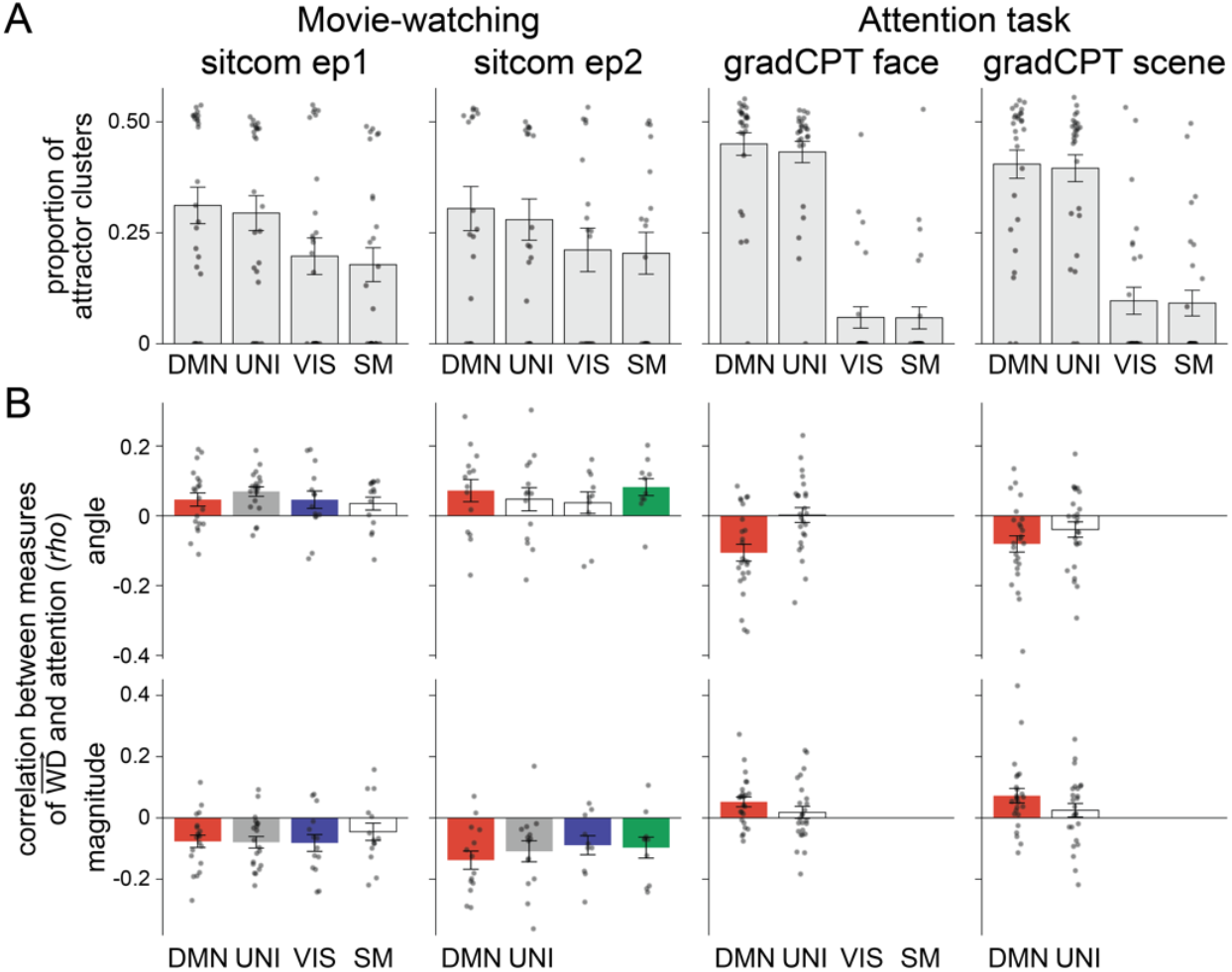
Relationships between attention dynamics and neural dynamics toward four attractors. The figure complements **Figure 5D** such that analyses were conducted separately for the two runs in each context.

## Footnotes

1 In dynamical systems and control theory, the intrinsic component is often referred to as autonomous, meaning its evolution depends on internal states, independent of external inputs.

2 Margulies et al. (2016) applied a nonlinear dimensionality reduction algorithm on hundreds of participants’ resting-state functional connectivity data to find top gradients that explained the largest variances. The primary gradient distinguished unimodal from transmodal areas, and the secondary gradient distinguished sensory from motor areas. These gradients were argued to be an “intrinsic coordinate system” of the human brain (Huntenburg et al., 2018) and has been replicated by multiple research groups (Bernhardt et al., 2022).

3 Narrative engagement has been characterized as a state of heightened emotional arousal and attentional focus (Busselle and Bilandzic, 2009; Bilandzic et al., 2019; Song et al., 2021; Ke et al., 2025). Because narrative engagement accompanies changes in both arousal and attentional states, denoting them as “attentional state” is a simplification made in this article.

